# Use-dependent biases primarily originate from a contaminated motor plan

**DOI:** 10.1101/2021.10.21.465112

**Authors:** Jonathan S Tsay, Hyosub E Kim, Arohi Saxena, Darius E Parvin, Timothy Verstynen, Richard B Ivry

## Abstract

Repetition of a specific movement biases subsequent actions towards the recently practiced movement, a phenomenon referred to as use-dependent learning (UDL). UDL has been attributed to shifts in the tuning of neurons in the motor cortex. However, recent studies employing a forced reaction time task, including the eLife article by Marinovic et al (2017), indicate that these biases may also arise from a contaminated motor plan, one that is biased towards the practiced direction. We advanced this line of inquiry, seeking to establish the relative contribution of execution and planning processes to UDL in a center-out reaching task in which participants were able to initiate movements of their own volition. On most trials, the target appeared at a designated “frequent” location; on other trials, the target appeared at one of six “rare” locations. In Experiment 1, participants exhibited a robust movement bias towards the frequent target when movements were self-initiated quickly, but a small movement bias when movements were self-initiated slowly – the signature of a contaminated motor plan. Strikingly, the heading angles were bimodally distributed, with one peak at the frequent target location and the other at the rare target location – a finding reinforced by a re-analysis of two widely cited studies on UDL. Notably, the latter peak was shifted in the frequently practiced direction, a signature of a motor execution bias. To eliminate the contribution of planning-related UDL, we imposed a delay between target onset and movement initiation in Experiment 2. As predicted, the heading angles became unimodally distributed around the rare target. The peak of this distribution was again shifted towards the location of the frequent target, indicative of a persistent bias in motor execution. Taken together, these results highlight two distinct components of UDL even when movements are self-initiated: First, the temporal dynamics underlying movement planning, in which a default plan is progressively overridden by a new plan, produces a pronounced motor planning bias. Second, there is a small, temporally stable bias that may reflect shifts in motor unit tuning.

## Introduction

Movement repetition can implicitly bias future movements to resemble the repeated movement (Classen, Liepert, Wise, Hallett, & Cohen, 1998), a phenomenon referred to as use-dependent learning (UDL). This similarity can be seen in features such as initial direction, speed, and magnitude of movements ranging from single-joint actions to whole-body locomotion (H. L. Chapman et al., 2010; Diedrichsen, White, Newman, & Lally, 2010; Hammerbeck, Yousif, Greenwood, Rothwell, & Diedrichsen, 2014; Huang, Haith, Mazzoni, & Krakauer, 2011; Jax, Buxbaum, & Moll, 2006; Jax & Rosenbaum, 2009; Wong, Goldsmith, Forrence, Haith, & Krakauer, 2017; Wood, Kim, French, Reisman, & Morton, 2020; Wood, Morton, & Kim, 2021). Theoretically, these movement biases have been attributed to shifts in motor unit tuning in the direction of a frequently practiced movement or potentiation of synapses that are frequently activated, resulting in long-term changes in movement execution (Butefisch et al., 2000; Celnik et al., 2006; Galea, Vazquez, Pasricha, de Xivry, & Celnik, 2011; Han, Arbib, & Schweighofer, 2008; Stefan, Classen, Celnik, & Cohen, 2008).

More recent work has upended this perspective, suggesting that movement biases primarily originate from limitations associated with motor planning. Specifically, when preparing to act, participants may first generate a default plan associated with a practiced or recent movement, then modify or override this plan when the context requires a different action. By this view, biasing effects should be most pronounced when planning time is limited. This hypothesis is supported by work in which the amount of time for planning was manipulated: Marinovic et al (2017) asked participants to make isometric forearm contractions to move a cursor to a visual target (Marinovic, Poh, de Rugy, & Carroll, 2017). Using a timed response task (Ghez, Hening, & Favilla, 1989), participants were provided with either a short (150 ms) or long interval (500 ms) to plan their movement towards frequently presented or rarely presented targets. Movement biases towards the frequently presented target were quite pronounced when preparation time was short, and biases were negligible when preparation time was long - a signature of a motor planning bias.

However, the countdown task employed by Marinovic et al may have unintentionally caused participants to adopt a default movement plan. Given that the instructions emphasized that movements must be initiated at the time of the imperative signal, it would be optimal for participants to anticipate that the target will appear at the frequent location, and thus, prepare a movement towards that location. When the target appears at a rare, unexpected location, the desire to initiate a default movement in synchrony with the imperative signal is in tension with the desire to re-aim towards the unexpected location. This tension should produce a planning bias, at least in the aggregate data: On some trials, the participant may initiate the movement when still in the default state; on other trials, they may have had sufficient time to replace the default plan, either partially or fully with a plan that is directed at the actual target.

We thus set out to examine UDL under conditions in which initiation time was unconstrained, and as such, allow planning to unfold in a more natural manner. On each trial, a target appeared at one of seven locations and the participant made a center-out reaching movement, attempting to intersect the target. To create the conditions for UDL, the target appeared at one location on 86.8% of the trials and the other locations on 2.2% of the trials each. We did not impose any constraint on initiation time. UDL, if present, should be evident in the distribution of the initial heading angle of the reaches. We assumed planning-related UDL would produce a broad distribution of heading angles between the frequent and rare target location. In contrast, execution-related UDL would manifest as a shift in the distribution of reaches towards the frequent target location. Moreover, an execution bias should be insensitive to planning time given the assumption that it arises from stable use-dependent plasticity involved in the implementation of motor commands.

Although the term “use-dependent” might suggest a form of Hebbian learning where the associative strength is only a function of repetition, evidence in both rodent (Dang et al., 2006; Verstynen & Sabes, 2011) and human work indicates that reward enhances use-dependent biases (Mawase, Lopez, Celnik, & Haith, 2018; Mawase, Uehara, Bastian, & Celnik, 2017a). Previous studies of UDL in humans have involved tasks in which reinforcement was always provided, either in the form of motivational verbal cues (Classen et al., 1998; Marinovic et al., 2017) or points to encourage faster movement initiation and/or movement accuracy (Mawase et al., 2018, 2017a; Verstynen & Sabes, 2011). These reinforcers, intended to enhance participants’ motivation, may have biased participants to adopt a default motor plan to maximize reward. We therefore eliminated any form of reinforcement in Experiment 1 and directly test the impact of reward on UDL in Experiment 2.

## Results

### Experiment 1

Participants performed center-out reaches, moving to a visual target that appeared at either a frequent location or rarely presented locations. We did not impose any constraint on reaction time, allowing the participants to initiate the movement at their own pace. To minimize on-line corrections, feedback was provided whenever the movement duration exceeded 400 ms. Since individual reaches were composed of straight and curved reaches (Fig S1), we examined the heading angle shortly after movement initiation, a time point that should index the participant’s initial movement plan (plus motor noise).

Movement biases towards the repeated location were evident in the reaches made to the probe locations, with all of the shifts towards the location of the frequent target (Fig 1B, inward bias). Averaging across participants and target direction (relative to the frequently repeated location), the size of the bias ranged from an average of 7.6° for the probe closest to the frequent target to 28.2° for the most distant probe location (main effect of probe distance: *χ*_(1)_ = 7909.3, *p* < 0.001, *η*^2^ = 0.5). Biases were negligible at the frequent target location (1.5° ± 0.7). When tested against the null, no-bias hypothesis, the biases for all of the probe locations were significant (±30°: 7.6 ± 2.1; *t*_9_ = 3.7, *p* = 0.005, *d* = 1.2; ±60°: 19.9 ± 4.2; *t*_9_ = 4.8, *p* < 0.001, *d* = 1.5; ±90°: 28.2 ± 6.7; *t*_9_ = 4.2, *p* = 0.002, *d* = 1.3). Moreover, the magnitude of the bias increased with probe distance (Bonferroni-corrected for three comparisons: ±30° vs ±60°: 12.3 ± 2.7; *t*_9_ = 5.2, *p* = 0.002, *d* = 1.6; ±60° vs ±90°: 8.2 ± 4.3; *t*_9_ = 2.2, *p* = 0.15, *d* = 0.7; ±30° vs ±90°: 20.5 ± 5.7; *t*_9_ = 4.2, *p* = 0.006, *d* = 1.3).

**Figure 1.**
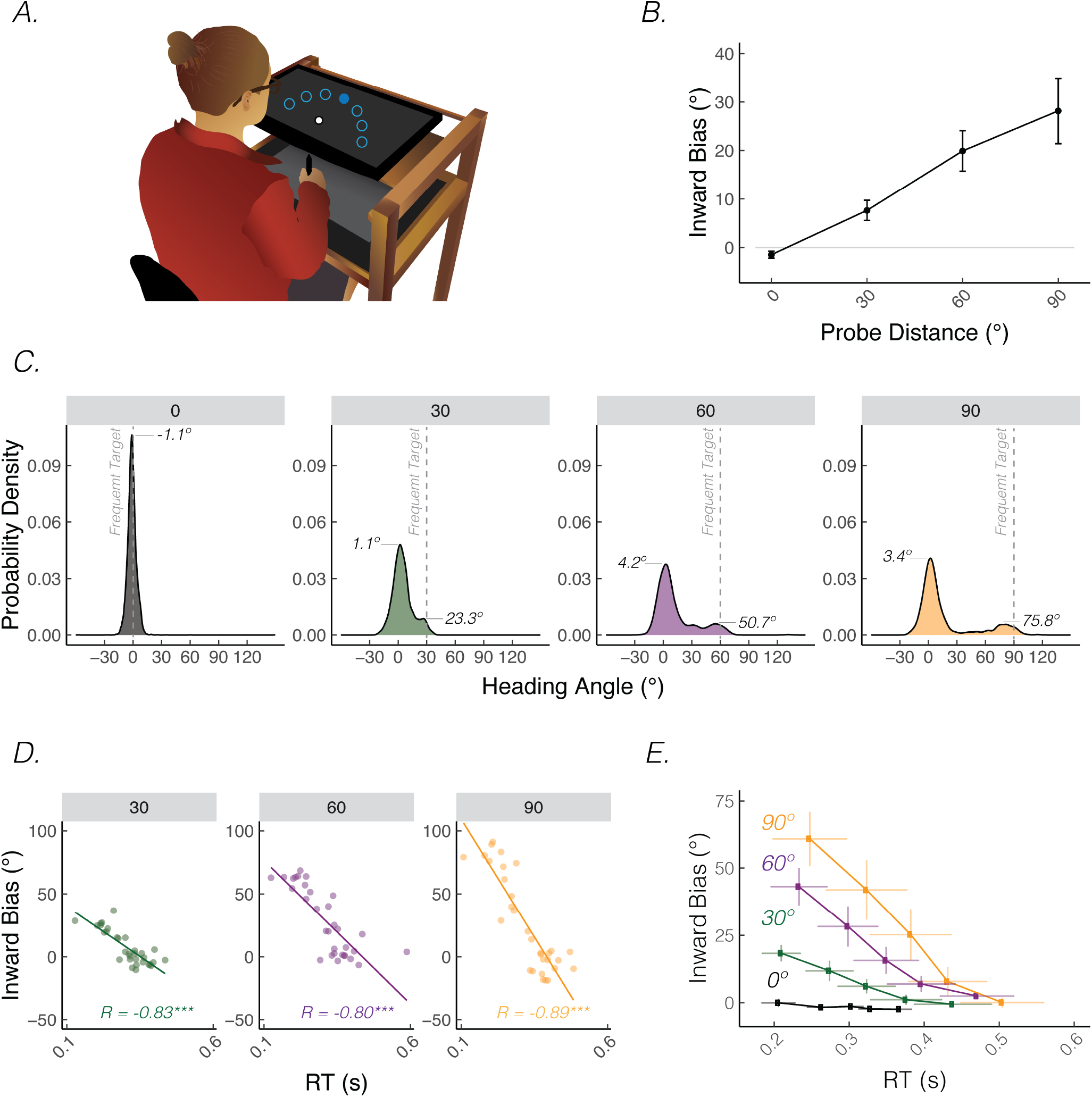
Hasty reaches elicited greater movement biases in Experiment 1. **A)** Reaching set-up showing locations of frequent and rare probe targets. Only one of seven targets (filled blue circle) was visible on each trial. **B)** Average inward biases increased as a function of probe distance. Data are pooled for the clockwise and counterclockwise pair that were equidistant to the frequent target location. **C)** Distribution of heading angles for each of the probe distances. Dashed line denotes the location of the frequently presented context target and 0 on the x-axis denotes the location of the probe target. The means obtained from the mixture of Gaussian model are provided. **D)** Bias as a function of a RT for a representative participant. Dots indicate individual reaches with the thin line showing the best fitting regression line. *R* denotes Pearson correlation; *** = p < 0.001. **E)** Group level analysis of bias as a function of RT. For each individual, RTs were binned into quintiles and mean bias was calculated for each quintile. These data were then averaged across the group. Error bars denote SEM.

We looked at the heading angle distributions to assess whether these mean biases reflected a shift in the heading angle distribution towards the mean bias, a mixture of reaches to the frequent and rare target locations, or a combination of these factors. We reasoned that use-dependent changes in tuning profiles should produce an overall shift in the distribution whereas a mixture of two types of movements would manifest as a bimodal distribution of heading angles. As shown in Fig 1C, the data for all probe distances are bimodal (likelihood ratio test of bimodal vs unimodal, probe distance 30°, 60°, 90°: all *p* < 0.001; probe distance 0° is unimodal: *p* = 0.08), with one peak at the probe location and a second peak near the frequent target location. In addition, the distributions include a wide range of intermediate heading angles. As such, the distribution of heading angles exhibits signatures of two distinct movement plans, one towards the frequent target location and one towards the rare target location, as well as an integration of these movement plans that results in some heading angles that are directed between these two locations. We return to this issue in the Discussion where we consider possible mechanisms that might account for action planning biases.

We next examined the relationship between heading angle and reaction time (RT). Given the bimodal nature of the heading distributions, we reasoned that faster reaction times would be associated with (incorrect) movements towards the frequent target location, whereas slower reaction times would be associated with (correct) movements towards the rare target. Consistent with this planning time hypothesis, we observed a strong negative relationship between heading deviation (bias) and reaction time. As can be seen in the data from one representative participant (Fig 1D), inward biases towards the frequent target were much larger when reaction time was fast compared to when reaction time was slow; indeed, there are a cluster of reaches to the frequent target location for the fastest RTs and a cluster of reaches to the rare probe target location for the slowest RTs.

This negative correlation was observed in the individual data for most participants (Fig S2) and at the group level for all of the probe distances (RT vs bias slope, 30°: −55.3 ± 3.1; *t*_7191_ = −17.8, *p* < 0.001; 60°: −99.1 ± 2.9; *t*_7191_ = −34.1, *p* < 0.001; 90°: −136.7 ± 2.5; *t*_7191_ = −5.7, *p* < 0.001). It was not observed for probe trials in which the target appeared at the frequent target location (0°: 1.2 ± 1.3; *t*_6965_ = 0.9, *p* = 0.35). To visualize this effect the RT data were segmented into five evenly sized bins (quintiles), with the bias data for each quintile averaged across participants (Fig 1E). To compare across the three rare probe locations, we normalized the bias data within each probe distance. The interaction was not significant (Fig S4, *χ*_(1)_ = 3.1, *p* = 0.21, *η*^2^ = 0.0), indicating that the RT-dependency was similar for all probe locations.

While the RT-based analyses suggest that a large component of use-dependent biases arises from limitations in action planning, there may nonetheless be a bias component that is associated with subtle changes in motor execution. The mean of the Gaussian centered around the rare location for each probe distance provides a rough estimate of a stable bias component, under the assumption that this inferred distribution is composed of reaches directed to the actual target. Consistent with the observed data, this value was shifted in direction of the frequent target location for each of the three probe distances (95% bootstrapped confidence interval from 10,000 samples is greater than 0°, the rare target location: 30°: [1.0°, 2.3°], 60°: [3.2°, 5.1°], 90°: [2.5°, 4.2°]). Thus, this analysis is consistent with the hypothesis that there is a small execution-related bias. We directly tackle this question in Exp 2, recognizing the limitations with this (mixture of Gaussian) model-based analysis.

The strong dependency of heading angle on RT, including the significant number of reaches that are erroneously directed to the frequent location, is somewhat puzzling given that there were no explicit instructions or constraints concerning RT. We speculate that participants may operate under a self-imposed urgency signal (Bogacz, Wagenmakers, Forstmann, & Nieuwenhuis, 2010; Murphy, Boonstra, & Nieuwenhuis, 2017). This could come about as spillover from the experimental instructions to “reach as quickly as possible” or some more generic factor such as a desire on the participants part to complete the experiment as fast as possible. Regardless of the underlying reason, the bimodal distribution of heading angles across all probe distances and the negative correlation seen with RT and bias indicate that limitations in motor planning underlie the large use-dependent effects observed in the current experiment.

### Reanalysis of Verstynen and Sabes (2011)

The results of Exp 1 motivated us to re-examine the data from Verstynen and Sabes (2011), a study frequently cited to demonstrate use-dependent effects on movement execution (despite the authors themselves being agnostic about whether biases arise from motor execution or planning). In particular, we were motivated to see if their data exhibited the three signatures of UDL observed in Exp 1: 1) Bimodality distribution of heading angles; 2) Shift towards frequent target location in distribution of reaches centered around the target; 3) Negative correlation between RT and bias. While the first two distributional features were also present in the data from Marinovic et al (2017) (Fig S5), the data were problematic for a correlational analysis relating RT and bias, since planning time was directly manipulated in their study. For this reason, we will focus on the data of Verstynen and Sabes, where planning time varied only with intrinsic factors.

The distribution of heading angles was bimodal at the 60° and 90° probe distances (likelihood ratio test of bimodal vs unimodal, probe distance 60°: *p* < 0.001, 90°: *p* < 0.001), with a clear peak at the rare target location and a smaller, second peak close to the frequent target location (Fig 2A). Bimodality was not discernable for the 30° probe distance (*p* = 0.86). As in Exp 1, there was a range of intermediate movements between the two peaks for these two larger probe distances. Thus, at these two probe distances, heading angles seem to again reflect a mixture of action planning to the frequent and rare probe location, as well as motor plans that result in movements with an initial heading between these two locations.

**Figure 2.**
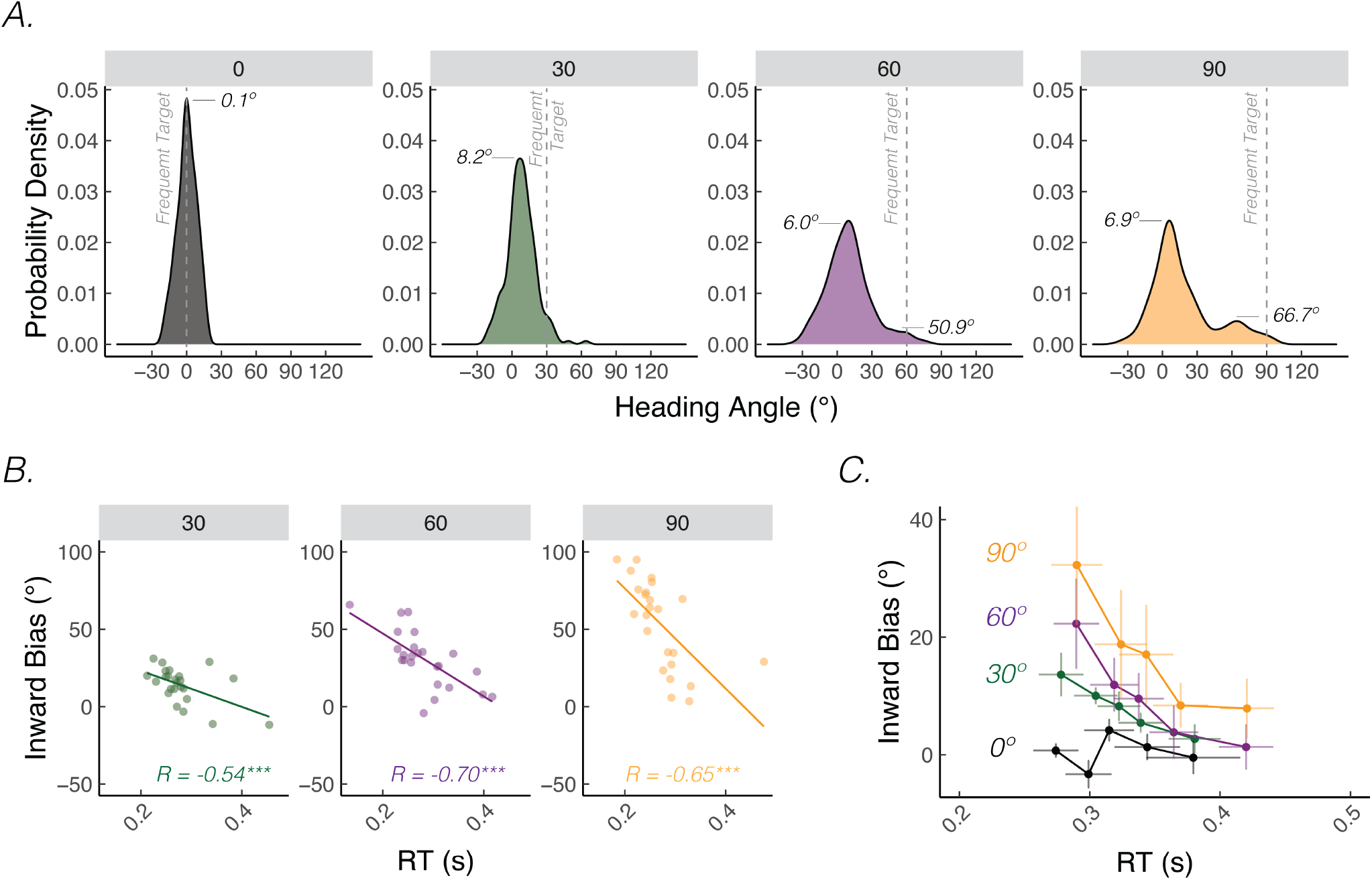
Hasty reaches elicited greater movement biases in Verstynen and Sabes (2011). **A)** Distribution of heading angles for each of the probe distances. Dashed line denotes the location of the frequently presented context target and 0 on the x-axis denotes the location of the probe target. The means obtained from the mixture of Gaussian model are provided. **B)** Representative individual’s bias vs RT function. *R* denotes Pearson correlation; *** denotes p < 0.001; Translucent dots denote individual reaches. Thin line denotes best fitting regression line. **C)** Group level quintile analysis of bias vs RT. Error bars denote SEM.

The magnitude of the biases in the Verstynen and Sabes data were also strongly dependent on reaction time. As seen in a representative individual (Fig 2B, S3 for all individuals) and in the group data (Fig 2C), movements initiated faster were associated with greater inward biases, an effect that persisted across all probe distances. The negative correlation between bias and RT was also robust on a group level for all probe distances (RT vs bias slope, 30°: -79.7 ± 21.3; *t*_642_ = −3.7, *p* < 0.001; 60°: −149.4 ± 17.9; *t*_648_ = −8.4, *p* < 0.001; 90°: −244.5 ± 17.3; *t*_645_ = −14.1, *p* < 0.001), other than 0°(−12.2 ± 22.8; *t*_660_ = −0.5, *p* = 0.59). When the bias data were normalized, there was no interaction between the three probe distances (Fig S4, *χ*_(1)_ = 0.8, *p* = 0.66, *η*^2^ = 0.0), indicating that the RT-dependency was similar for all probe locations.

We again focused on the peak around the rare target locations to obtain a rough estimate of a stable bias component. Similar to Experiment 1, the mean of these Gaussians was shifted in the direction of the frequent target location (Fig 2A), with a confidence interval that excluded zero for all three rare probe locations (95% bootstrapped confidence interval from 10,000 samples: 30°: [6.8°, 9.8°], 60°: [3.7°, 8.6°], 90°: [4.5°, 9.3°]).

In summary, our re-analysis of the data from the Verstynen and Sabes (2011) points to the presence of two forms of UDL, similar to that observed in Exp 1. The large bias inferred from the aggregate data is likely driven by limitations in action planning. As shown in the correlational analysis, planning-related biases are closely related to RT. In addition, there is a small shift in the distribution of reaches directed close to the rare target location, perhaps originating from a bias in motor execution. We assume that execution-related biases, if reflecting a structural change in motor unit tuning, would be independent of RT. We test this idea in the second experiment.

### Experiment 2

The results of Exp 1 and our re-analysis of the Verstynen and Sabes (2011) data indicate that a large component of use-dependent bias is related to limitations in action planning. We also observed a small shift in distribution of heading angles (correctly) directed towards the actual target, and hypothesize that this pattern of results arises from a small use-dependent movement execution bias. Our analysis of this component was indirect, inferred by the peak around the probe location in the mixture of Gaussians model. As a more direct test, we opted to use a delayed-response task in Exp 2, imposing a 500 ms delay between the presentation of the target location and imperative signal. We reasoned that the additional preparation time would allow participants to plan their actions in advance of the imperative, minimizing RT-dependent effects on movement biases while providing a more sensitive measure of use-dependent biases in movement execution.

We also used Exp 2 to examine the influence of reward on execution-related UDL, by comparing a group who received binary, rewarding feedback following accurate movements to the frequent target (Reward group) to a group who received no feedback (No Reward group). Importantly, the No Reward group represents a “pure” use-dependent condition as they never received any type of feedback regarding task success, including any potential intrinsic reward from seeing a cursor hit the target (Kim, Parvin, & Ivry, 2019; Tsay, Haith, Ivry, & Kim, 2021).

We first asked if biases persist when planning time is extended. The distributional analysis showed a marked difference than that observed in Exp 1. The distribution of heading angles for all probe distances (in both groups) were tightly clustered around the rare target location, with minimal heading angles in the direction of the frequent target location (Fig 3A). Indeed, the distributions were all best described as unimodal (likelihood ratio test of bimodal vs unimodal, all probe distances: *p* = 1), with a single peak near the location (rare or frequent) at which the target appeared. Although the delay yielded unimodal distributions, the peak of the distributions was not centered at the probe locations but instead consistently shifted in the direction of the frequent target location (95% bootstrapped confidence interval from 10,000 samples: 30°: [3.0°, 3.7°], 60°: [3.3°, 3.9°], 90°: [2.5°, 3.2°]).

**Figure 3.**
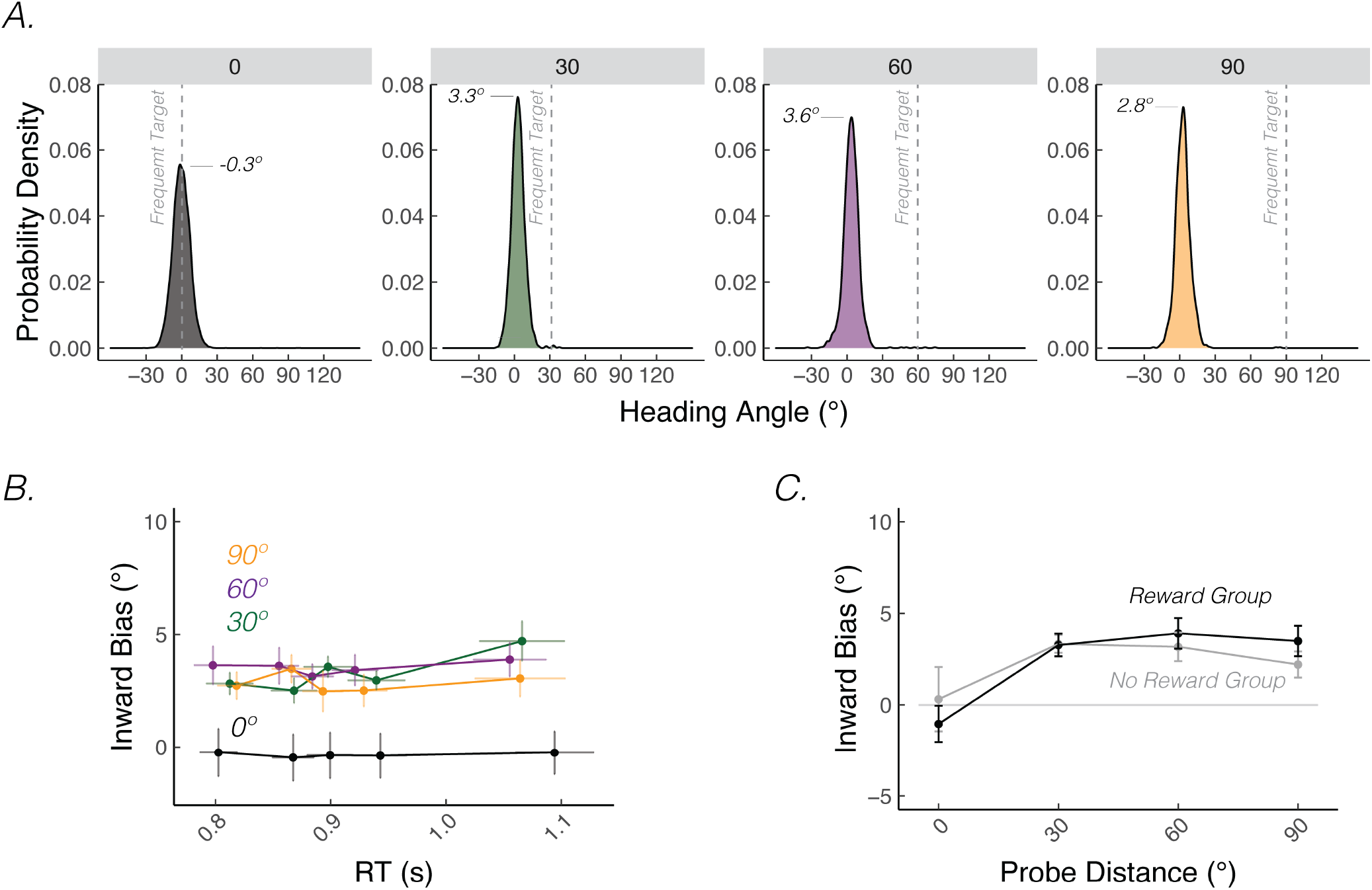
Use-dependent motor execution biases were small and not modulated by reward in Experiment 2. **A)** Distribution of heading angles for each of the probe distances. Dashed line denotes the location of the frequently presented context target and 0 on the x-axis denotes the location of the probe target. The means obtained from the mixture of Gaussian model are provided. **B)** Group level quintile analysis of bias vs RT. Reward and No Reward groups were combined in this panel since RTs and movement biases, since both variables did not vary with reward feedback. Error bars denote SEM. **C)** Average inward biases were modest for both Reward and No Reward groups.

This persistent, albeit small bias towards the frequent target for all probe distances was corroborated by a series of post-hoc t-tests (30°: 3.3 ± 0.4; *t*_31_ = 8.6, *p* < 0.001, *d* = 1.5; 60°: 3.5 ± 0.6; *t*_31_ = 6.2, *p* = 0.001, *d* = 1.1; 90°: 2.8 ± 0.6; *t*_31_ = 5.2, *p* < 0.001, *d* = 0.9). The bias was not present for probe reaches to the frequent target location (0°: −0.4 ± 0.6; *t*_31_ = −0.4, *p* = 0.72, *d* = 0.1). The magnitude of the bias did not increase with probe distance (Bonferroni corrected for three comparisons: ±30° vs ±60°: 0.2 ± 0.5; *t*_31_ = 0.5, *p* = 1, *d* = 0.1; ±60° vs ±90°: 0.7 ± 0.4; *t*_31_ = 1.6, *p* = 0.33, *d* = 0.3; ±30° vs ±90°: 0.5 ± 0.5; *t*_31_ = 0.9, *p* = 1, *d* = 0.2). Thus, the results indicate that with extended planning time, a small use-dependent bias is observed.

Unlike Exp 1, the magnitude of the bias in Exp 2 was not negatively correlated with RT for any of the probe distances (Fig 3B) (RT vs bias slope, 30°: 4.2 ± 1.1; *t*_24726_ = 3.7, *p* < 0.001; 60°: 1.6 ± 1.3, *t*_24723_ = 1.2, *p* = 0.24; 90°: −1.6 ± 0.9, *t*_24723_ = −1.7, *p* = 0.09). Curiously, there was a small positive correlation for probe distance 30°, with the bias slightly larger on trials with longer RTs (Fig 3B). While the cause of the positive slope is unclear, the effect was modest (4.2°/s) in comparison to values observed in Experiment 1 and Verstynen and Sabes data (range between -55.3°/s and -244.5°/s).

We next examined the influence of reward on the small biases observed across probe targets, asking first if the reward manipulation was effective. To this end, we compared movement accuracy between the two groups for reaches to the frequent target (the only location eligible for reward in the Reward group). As shown in the cumulative distribution functions (Fig S6), participants in the Reward group were more accurate than those in the No Reward group (*t*_30_ = 3.2, *p* = 0.003, *d* = 1.1): That is, reward provided at the target location was more likely to produce reaches with smaller absolute angular deviation from the target. As such, our reward manipulation had an appreciable effect on behavior. However, the reward manipulation had no effect on bias (Fig 3C) (absent main effect of reward: 0.1 ± 0.9; *χ*_(1)_ = 0, *p* = 0.92, *η*^2^ = 0.02; absent reward x probe distance interaction: *χ*_(1)_ = 4.3, *p* = 0.12, *η*^2^ = 0).

Taken together, the results of Exp 2 highlight three points. First, the large biases towards a frequent target, the signature of use-dependent learning, were largely abolished by the introduction of a short delay period between the presentation of the target and the imperative signal. Although the inference is based on a between-experiment comparison, this finding is consistent with the hypothesis that use-dependent bias in reaching largely arises from limitations in movement planning. Second, the small residual bias may reflect use-dependent changes in processes associated with movement execution. Third, this bias was not modulated by reward, challenging the popular belief that use-dependent and reinforcement learning processes interact.

## Discussion

Sensorimotor learning entails multiple learning processes (Kim, Avraham, & Ivry, 2020; Krakauer, Hadjiosif, Xu, Wong, & Haith, 2019; Shadmehr, Smith, & Krakauer, 2010), including mechanisms sensitive to errors, rewards, and strategy use. Use-dependent learning refers to behavioral changes that arise from repetition. Classically, use-dependent changes have been attributed to changes in representations associated with motor execution (e.g., tuning functions of output units) (Classen et al., 1998). TMS studies provided some of the seminal evidence in favor of this hypothesis, where muscle-activation patterns in response to single-pulse suprathreshold TMS dramatically biased movements toward the axis of a recently practiced single-joint movement (Classen et al., 1998; Flöel et al., 2005). Given that the TMS pulses directly activated the corticospinal tract, the authors concluded that use-dependent learning reflected plasticity in motor cortex. A similar interpretation has been proposed based on use-dependent effects observed following repeated passive movement in a given direction (Diedrichsen et al., 2010; Javidialsaadi, Albert, & Wang, 2021).

Recent work has shown that use-dependent learning may also reflect biases associated with action planning. For example, the plan required to move to a rare target may be contaminated by a plan associated with a frequent target. This may be due to residual activation of a recent action plan or an expectancy bias for the frequent target. In support of the planning hypothesis, Marinovic et al (2017) showed that use-dependent biases during wrist flexion/extension were markedly reduced when planning time was increased from 150 ms to 500 ms. A similar effect of preparation time was also observed in eye movements (Reuter, Marinovic, Welsh, & Carroll, 2019); indeed, under short preparation time, participants occasionally made a saccade to the frequent target location prior to the onset of the actual target.

We set out to ask whether dissociable components of UDL would manifest even in a more naturalistic (unpressured) environment, where participants are free to initiate their movements at their own volition. We also eliminated any reinforcing feedback in Experiment 1 to tightly control for any potential confounding effects of reward (Flöel et al., 2005). When cued to reach to rare target locations, participants exhibited a bias towards the frequent target location, and the magnitude of this bias increased with probe distance. We found that this bias was strongly correlated with reaction time: Participants exhibited larger biases with short reaction time, and negligible biases when preparation time was long – a finding that consistent with the proposal by Marinovic et al. (2017). However, across all conditions reported here, we also observed a small bias towards the frequent target location that was independent of RT. Strikingly, this temporally-stable bias was consistently seen in the distribution of heading angles, whereby the peak of the distribution located near the rare probe target was shifted towards the frequent target. We hypothesize that this bias is the mark of a true use-dependent bias in motor execution.

### Potential mechanisms that account for motor planning biases

We consider here three hypotheses that may account for how use-dependent biases come about from constraints on motor planning: the switching hypothesis (two variants: categorical vs gradual switching), the performance optimization hypothesis, and the mental transformation hypothesis. First, the switching hypothesis is based on the idea that the planning system maintains a representation of a default plan directed towards the frequent target location, and that this default plan must be replaced when the target appears elsewhere. Use-dependent biases arise from the dynamics associated with this transition. At one extreme, the transition may be a categorical process — at some point the system undergoes a phase transition such that the new plan abruptly replaces the default plan (Wong, Haith, & Krakauer, 2015). This model would predict a bimodal distribution, with an initial plan directed towards the frequent target location abruptly overridden by a plan towards the probed target. This bimodality was observed in the distributional analyses of Exp 1 and Verstynen and Sabes.

However, in the other extreme, other aspects of the data suggest that the switching process may entail a more gradual transition. The heading angles spanned a wide range of intermediate movements between the rare and frequent target locations. It seems unlikely that these intermediate heading directions could result from motor noise; for example, an initial heading direction that is deviated 45° from the actual target. A gradual process would presumably yield a movement plan that is the weighted average of the decaying default plan and emerging target-defined plan. There is considerable behavioral evidence in support of this view (C. S. Chapman et al., 2010; Enachescu, Schrater, Schaal, & Christopoulos, 2021; Gallivan, Stewart, Baugh, Wolpert, & Flanagan, 2017; Stewart, Gallivan, Baugh, & Flanagan, 2014). Moreover, single neuron activity in premotor cortex has been hypothesized to reflect overlapping activity of two motor plans (P. Cisek & Scott, 1999; Paul Cisek, 2006; Paul Cisek & Kalaska, 2005) that eventually converges to one plan corresponding to the movement direction. A gradual switch could yield both a bimodal distribution (strong weighting for one plan or the other), as well as intermediate heading angles.

Whereas the switching hypothesis is based on the idea that there are distinct representations for movement plans to the probed targets and the frequent target location, the other two hypotheses posit a unitary representation. One variant is the performance optimization hypothesis, motivated by evidence suggesting that a default plan is flexibly optimized to minimize some task-dependent cost function (Alhussein & Smith, 2021; Dekleva, Ramkumar, Wanda, Kording, & Miller, 2016; Haith, Huberdeau, & Krakauer, 2015). For example, Haith et al. (2015) observed that, when planning time was limited, participants’ movements were directed towards intermediate locations when the potential targets were close to each other and bimodally distributed when the potential targets were far apart. A switching mechanism cannot readily accommodate this spacing interaction. Rather, the authors propose that when potential targets were close, a default plan to an intermediate position is optimal because an on-line adjustment may suffice to reach the target; however, when potential targets are far, such online adjustments are unlikely to reach the target, and thus the optimal plan is to anticipate one location or the other. The distribution of heading directions in Exp 1 and Verstynen and Sabes (2011), while anecdotal, indeed appear more bimodal as the probe distance increased.

The other variant in which planning entails a unitary representation is the mental transformation hypothesis. In this view, there is a default movement plan that is transformed in a continuous manner (e.g., rotated) when the target appears at an unexpected location. For example, in the sensorimotor adaptation literature, strategic aiming appears to entail such a rotation process, one that starts at the target location and terminates at the desired aiming location (Anguera, Reuter-Lorenz, Willingham, & Seidler, 2010; McDougle & Taylor, 2019; Pellizzer & Georgopoulos, 1993). Population vectors in the motor cortex show evidence of such a rotational process when the animals are required to reach to a location that requires a spatial transformation (Georgopoulos, Lurito, Petrides, Schwartz, & Massey, 1989). By this view, the distribution of heading directions will be a function of the interaction of the speed of the rotation process and an internal urgency signal.

The current data do not allow us to distinguish between these three hypotheses concerning the marked use-dependent bias associated with motor planning. All can account for the bimodal distributions, the trials yielding intermediate heading angles, and the strong dependency between RT and bias. We expect that it may be difficult to evaluate and compare the merits of these three hypotheses from behavioral data alone. However, complementary physiological measures may offer more definitive signatures supporting one hypothesis over the others; for example, neural population dynamics would likely diverge when based on the integration of two movements plans or optimization and/or transformation of a unitary movement plan (Dekleva, Kording, & Miller, 2018; Zimnik & Churchland, 2021).

### Biases that arise from use-dependent changes in motor execution

The RT analysis in Exp 1 revealed the transitory nature of use-dependent biases, an observation that underscores the argument that a major source of bias arises from dynamic changes occurring during motor planning. This interpretation is further supported by the results of Exp 2 where the insertion of a delay between the onset of the target and movement initiation eliminated much of the bias. Nonetheless, the data also reveal a second source of use-dependent bias, one that is temporally stable (i.e., independent of RT) and not related to the angular difference between the probe location and the frequent target location. While this bias might arise from some persistent attractor in planning space, we hypothesize that the distance and time independence are signatures of a bias that reflects a use-dependent change associated with motor execution.

Where we obtained a relatively “pure” measure of this motor execution bias in Exp 2, one uncontaminated by the dynamics of motor planning, we only observed a small bias. This value is consistent with the magnitude observed in our indirect assay of use-dependent movement execution biases in Exp 1 and reanalysis of the data of Verstynen and Sabes (2011): Across the different conditions, the mean of the Gaussian centered around the probe target was shifted by 6.6° (see Figs 1C and 2A). A similar estimate of residual bias was reported by Marinovic et al (2017) when preparation time was lengthened (Fig S5). Perhaps even more compelling evidence that this bias is related to processes associated with movement execution comes from Exp 2 of Diedrichsen et al (2010): In that experiment, use-dependent effects were tested by having participants make reaching movements following a set of passive limb displacements. In one condition the passive movements terminated at the central location of a visual target; in another condition, the passive movement deviated from the target by 8°. The latter resulted in a reaching bias of 2° in the direction of the passive displacement. Given that participants were unaware of the difference in the passive trajectories between the two conditions, it seems unlikely that this effect would be due to a use-dependent effect in motor planning.

Previous studies have indicated that reward can enhance plasticity effects in the motor cortex (Hollerman & Schultz, 1998; Luft & Schwarz, 2009; Wise, 2004). For example, in humans, a single oral dose of levodopa (precursor to dopamine) has been shown to increase the propensity in which movements elicited by suprathreshold TMS are biased in the direction of a recently practiced movement (Flöel et al., 2005) (see also, (Mawase, Uehara, Bastian, & Celnik, 2017b)). However, we failed to observe any effect of reinforcement on the magnitude of the motor execution bias in Experiment 2: The small use-dependent effect was similar for participants who received positive feedback following successful reaches as for those who did not receive any feedback (including visual), despite clear evidence that the group receiving positive feedback performed better at the task. At present it is unclear why our null results diverge from those observed in other studies.

Mechanistically, these use-dependent motor execution biases are attributed to a stable change in the sensorimotor map, such that the output in response to a given motor command is altered by experience. For use-dependent learning, this change might come about from a Hebbian-like mechanism where the weights in the map are altered in the direction of the practiced movement. Similarly, in error-based learning (e.g., sensorimotor adaptation), the weights are altered such that the same motor command will result in an output that should reduce the error. Critically, in both learning processes, behavioral changes attributed to alterations to the sensorimotor map are limited in magnitude (Avraham, Ryan Morehead, Kim, & Ivry, 2021; Bond & Taylor, 2015; Kim, Morehead, Parvin, Moazzezi, & Ivry, 2018; Morehead, Taylor, Parvin, & Ivry, 2017; Tsay, Avraham, et al., 2020; Tsay, Ivry, Lee, & Avraham, 2021; Tsay, Kim, Parvin, Stover, & Ivry, 2021; Tsay, Parvin, & Ivry, 2020); instead, behavior changes are primarily attributed to changes in motor planning (e.g., strategic re-aiming) (McDougle, Ivry, & Taylor, 2016).

This last point is relevant when considering the contribution of use-dependent learning to skill acquisition. The behavioral changes associated with skill acquisition have been hypothesized to reflect long-term reorganization of primary motor cortex following extended practice. For example, dramatic improvements in sequence production have been shown to be associated with increased levels of activation in primary motor cortex (Karni et al., 1995). However, recent work by Berlot et al (2020) failed to find evidence of sequence representation within primary motor cortex, or even changes in overall activation as a function of practice. Similar to the results of the current study, the benefits of long-term practice may reflect flexible, upstream processes involved in motor planning, rather than the small use-dependent changes associated with motor execution.

## Methods

### Participants

A total of 42 participants (mean age = 20 ± 2.2 years) were recruited for two experiments. The sample sizes were based on similar reaching studies assessing use-dependent learning (Marinovic et al., 2017; Verstynen & Sabes, 2011). All participants were right-handed as verified with the Edinburgh Handedness Inventory (Oldfield, 1971), and received course credit or financial compensation for their participation. The experimental protocol was approved by the Institutional Review Board at the University of California, Berkeley.

### Reaching Task

The participant was seated at a custom-made table that housed a horizontally mounted LCD screen (53.2 cm by 30 cm, ASUS), positioned 27 cm above a digitizing tablet (49.3 cm by 32.7 cm, Intuos 4XL; Wacom, Vancouver, WA) (Fig 1A). Stimuli were projected onto the LCD screen. The experimental software was custom written in Matlab using the Psychtoolbox extensions (Brainard, 1997).

Participants made center out reaching movements, sliding a modified air hockey paddle containing an embedded digitizing stylus across the tablet. The tablet recorded the position of the stylus at 200 Hz. Vision of the hand was blocked by the monitor, and the lights in the room were turned off to block peripheral vision of the arm.

At the beginning of each trial, participants moved their right hand to position the digitizing pen within a “start” circle (0.6 cm diameter open white circle in the center of the LCD screen). To assist the participant in finding this starting position, a white feedback cursor (0.5 cm diameter) appeared when the hand was within 2 cm of the start circle. The position of the cursor was aligned with the digitizing pen. Once the pen remained within the start circle for 500 ms, the target appeared (blue circle, 1 cm diameter). The radial position of the target was always 10 cm from the start circle. In terms of angular position, the target could appear at one of seven locations, 0°, 30°, 60°, 90°, 120°, 150°, and 330° (Fig 1A). Participants were instructed to reach towards the target, making the movement in a smooth, rapid manner. To discourage online corrections, participants were told to slice through the target rather than attempt to stop at the target. A trial ended when the reach amplitude exceeded 10 cm or when the movement time exceeded 400 ms. Cursor feedback was presented throughout the first 10 cm of the movement trajectory and then remained visible at the target radius for 50 ms before turning off.

### Experiment 1

Ten participants were tested in Experiment 1. To create the conditions for use-dependent learning, the target appeared at one location with a much higher probability than at the other six locations (86.8% vs 2.2% for each of the other six locations). For half of the participants, the frequent target location was 60°; for the other half of the participants, the frequent target location was at 150°.

Each participant completed two baseline blocks and eight test blocks. In the first baseline block, the target appeared ten times at each of the seven locations. Cursor feedback was presented during the reach. The second baseline block consisted of another 10 reaches to the seven target locations, but no feedback was presented during the reach (or at the endpoint), allowing an estimate of each participant’s idiosyncratic biases.

The main experiment consisted of eight test blocks of 90 trials each. Following the design of Verstynen and Sabes (2011), each block started with 10 reaches to a target appearing at the high probability location to clearly establish this as the frequent target. This was followed by 80 more trials. Of these, the target appeared at the frequent location on 66 trials and feedback was provided during the reach. For the other 14 trials, the target appeared at one of the seven locations (including the frequent location) and the reaches were made without feedback. We did not provide feedback on these 14 probe trials to ensure that the participant was unaware of his or her bias. The order of the last 80 trials in each block was pseudorandomized, such that there was one probe trial every seven reaches.

The instructions emphasized that the reaches should be made quickly and in one smooth motion, attempting to intersect the target. To discourage movement speeds that might allow for online corrections, the message “too slow” was played over the computer speaker when movement time was greater than 400 ms. We did not place any emphasis on reaction time; participants were free to set their own pace for movement initiation. The error message, “too fast” was played over the computer speaker if the participant initiated the within 70 ms of target onset, a criterion set to eliminate anticipatory movements.

### Experiment 2

The general procedure was the same as in Exp 1, with three exceptions. First, we modified the session structure in Exp 2. The experiment began with the same two baseline blocks, one with online cursor feedback (70 trials) and one with no feedback (70 trials). This was followed by six test blocks of 134 trials. Within each test block, there were 113 reaches to the frequent target with potential reward feedback (see below) provided and 21 reaches to all seven locations with no feedback provided (3 trials/target, probe trials).

Second, we imposed a 500 ms delay between target appearance and an imperative cue, a tone played over the computer loudspeaker. By providing a 500 ms interval to prepare the movement, we sought to reduce or eliminate temporal constraint (either experimenter- or participant-imposed) on action planning.

Third, we eliminated the online cursor feedback (other than during the first baseline block) so that we could examine the impact of reward on movement biases. Participants were randomly assigned to one of the two groups (n=16/group), a Reward group and a No Reward group. For the Reward group, reaches to the frequent target were rewarded when the hand angle at the target amplitude was within ± 5.7° of the target. On these trials, the target turned green, doubled in size (to 2 cm in diameter), and a pleasant “ding” was played. If the hand angle was greater than ± 5.7° of the frequent target, no visual or auditory feedback was provided. For the No Reward group, no feedback was provided on any of the reaches.

### Data Analysis

Hand angle was defined as the angle between a line from the start position to the target and a line from the start position to the hand position, measured 50 ms after movement onset. By taking the estimate at 50 ms, we should eliminate any contamination from on-line corrections.

We considered two types of bias. First there is the bias to reach to a given target independent of the effects of the experimental manipulation. For each target location, we determined the participant’s baseline bias as the mean angular deviation during the baseline no-feedback block. This value should reflect any bias associated with regression to the mean (i.e., reaches to the center of the workspace) rather than biases due to repeated reaches to a frequently presented target. This value was thus subtracted from each reach to the corresponding target (both frequent and rare targets) in the training block. Second, to calculate use-dependent bias, the frequent target (60° or 150°) was reset to 0° and the other six targets were defined with respect to the frequent target (±30°, 60°, or 90°). The sign of the biases was flipped, such that positive values corresponded to biases towards the frequent target (i.e., inward bias) and negative values corresponded to biases away from the frequent target.

These biases were evaluated using a linear mixed effect model (R: lmer function) with RT and probe target distance as fixed (interacting) factors and participant as a random factor (Kuznetsova, Brockhoff, & Christensen, 2017). Satterthwaite’s degrees of freedom was also provided (Satterthwaite, 1946). Post-hoc t-tests on the betas from the linear mixed effect model (i.e., main effect of RT, probe distance, and interaction of RT x probe distance) were evaluated using emmeans and anova functions in R. The distribution of these biases (heading angles) was also modeled using a mixture of either one or two Gaussians (R: Mclust). A likelihood ratio test was used to discern which of these two distributions provided a better fit to the distribution (R: mclustBootstrapLRT).

Reaction time (RT) was defined as the position of the hand when hand velocity exceeded 3 cm/s. We binned each participant’s RTs into quintiles, with the first quintile composed of the fastest 20% of reaches and the fifth quintile composed of the slowest 20% of reaches. For each quintile, we calculated the mean bias. The mean RT and bias data for each quintile were then averaged across participants.

All post-hoc t-tests were two-tailed, and Bonferroni corrected for multiple comparisons. Standard effect sizes are reported (*η*^2^ for fixed factors; Cohen’s *d*_*z*_ for within-subjects t-tests, Cohen’s *d* for between-subjects t-tests, Pearson correlation R for linear regression) (Lakens, 2013).

## Supplemental Figures

**Figure S1.**
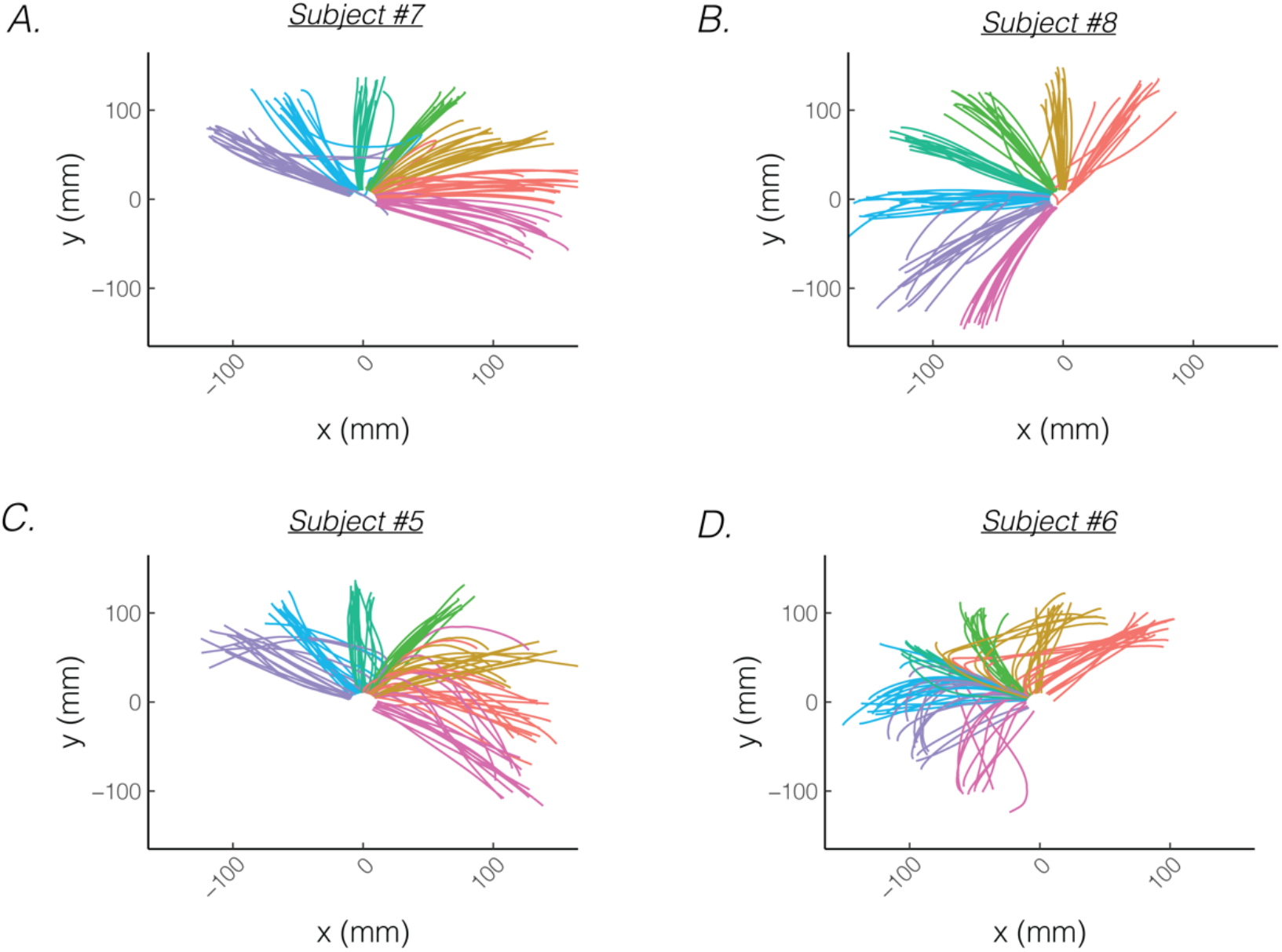
Reaches consist of a mixture of straight and curved movements in Experiment 1. Biases from four representative participants who exhibit primarily straight **(A, B)** vs curved movements **(C, D)**. The frequent target is denoted by the green curve, whereas rare targets are denoted by the purple, blue, turquoise, yellow, red, and magenta curves (clockwise).

**Figure S2.**
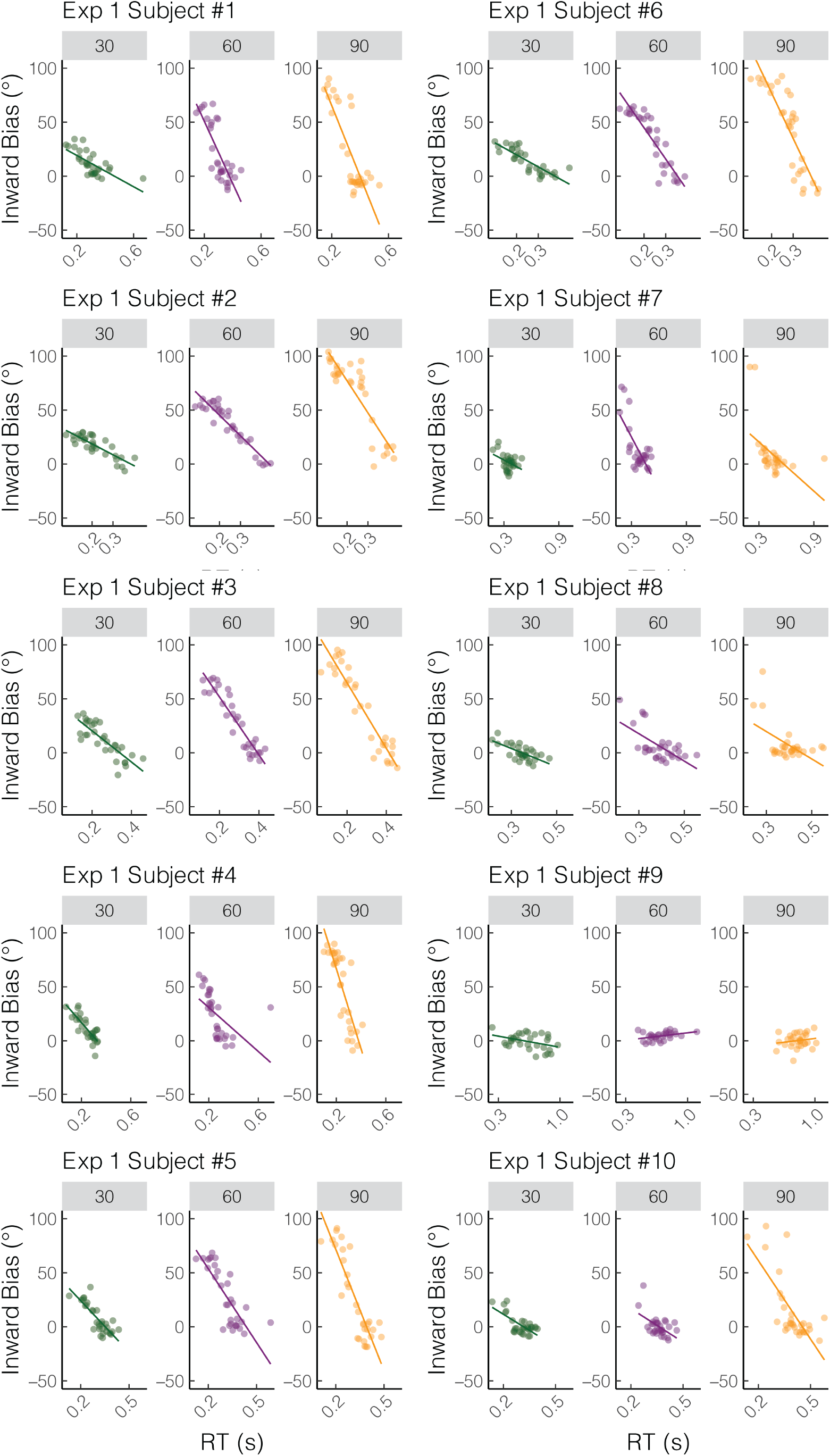
All individuals from Experiment 1.

**Figure S3.**
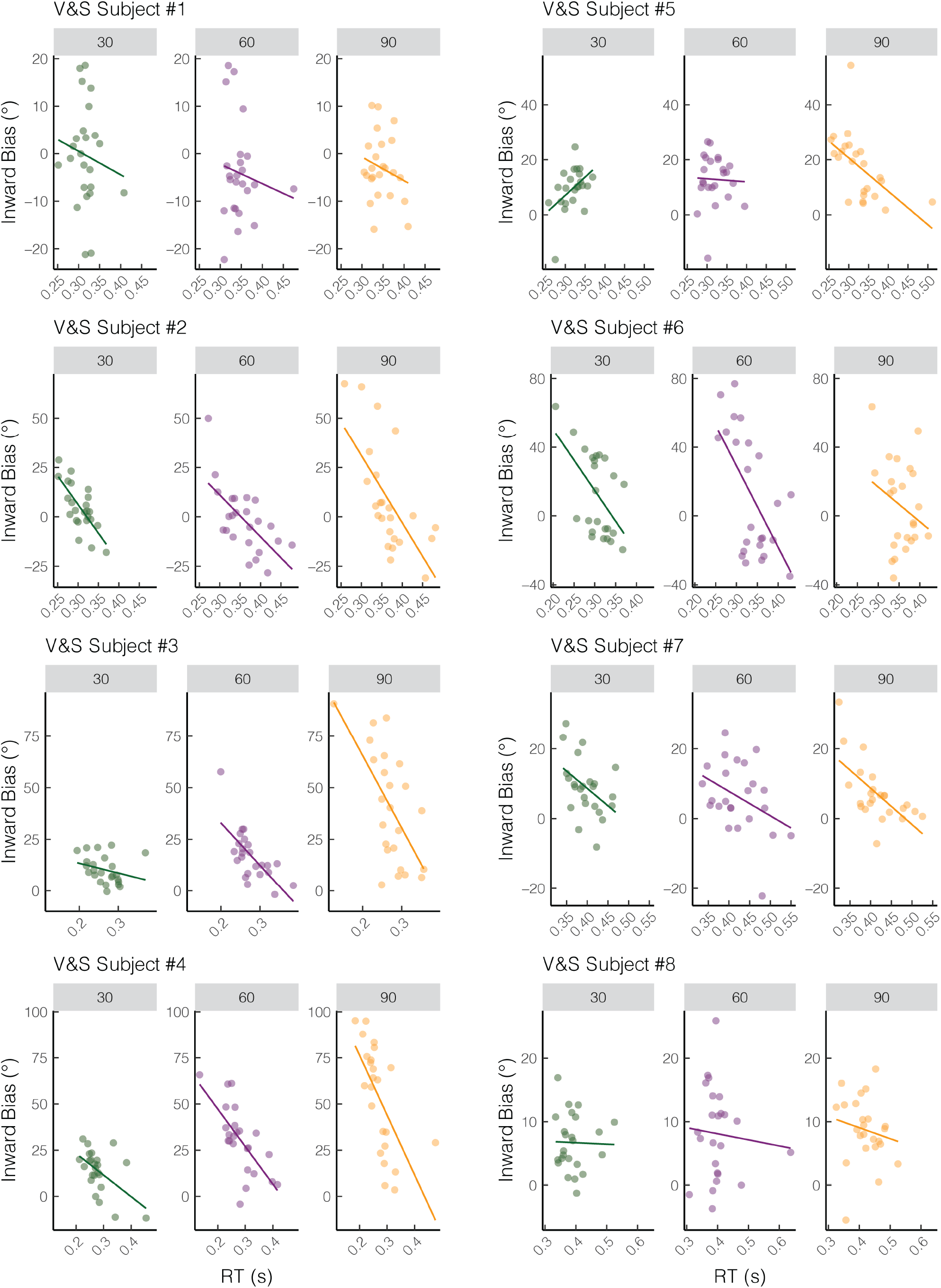
All individuals from Verstynen and Sabes (2011).

**Figure S4.**
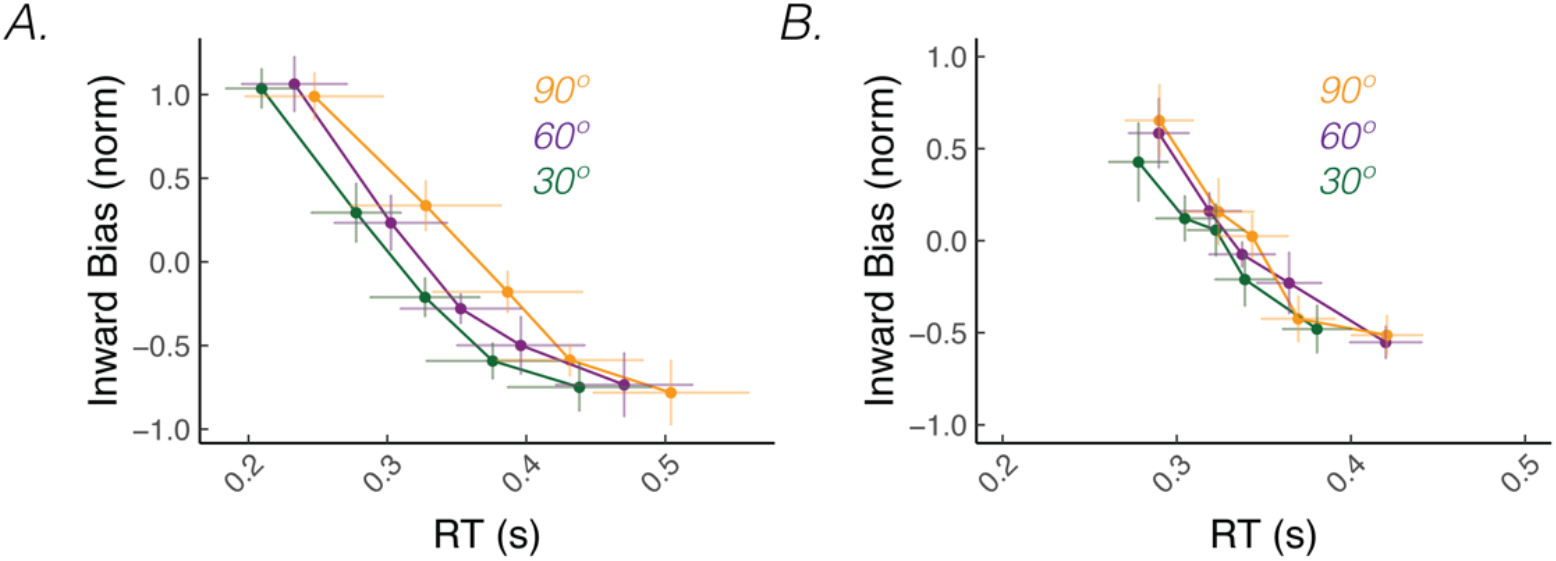
Biases from different probe distances have a similar dependency on RT. **A)** Group level quintile analysis of bias vs RT for Experiment 1 and **B)** Verstynen and Sabes (2011). Biases were normalized within each probe distance. Error bars denote SEM.

**Figure S5.**
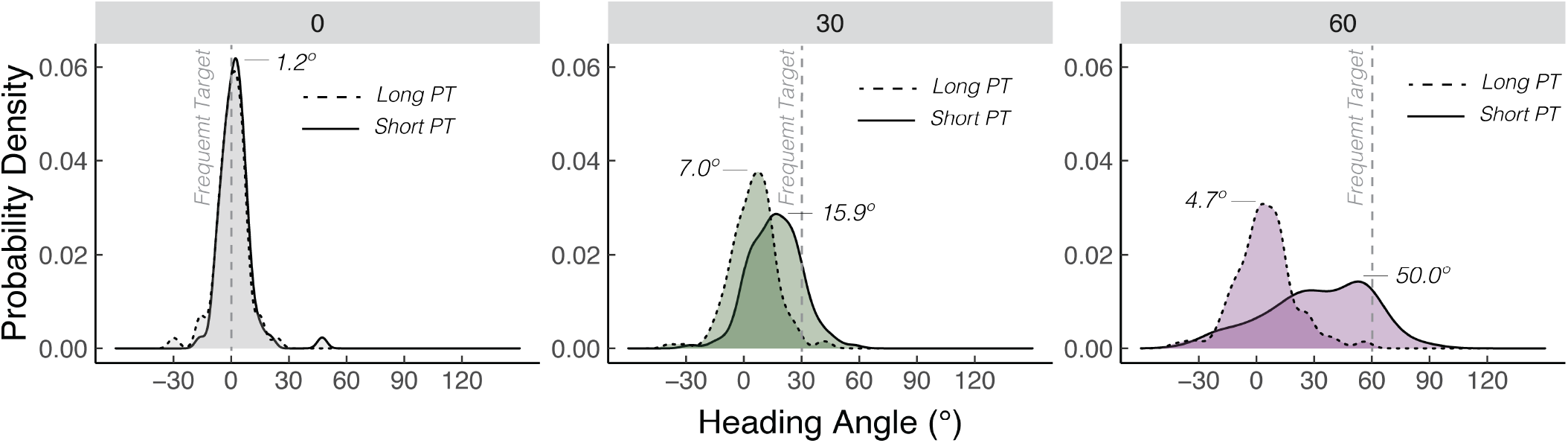
Distribution of heading angles across three probe distances in Marinovic et al (2017). Dashed vertical denotes the location of the frequently presented context target and 0 on the x-axis denotes the location of the probe target. Dashed curve denotes wrist movements with long preparation time and solid curve denotes wrist movements with short preparation time. The means obtained from the mixture of Gaussian model are provided.

**Figure S6.**
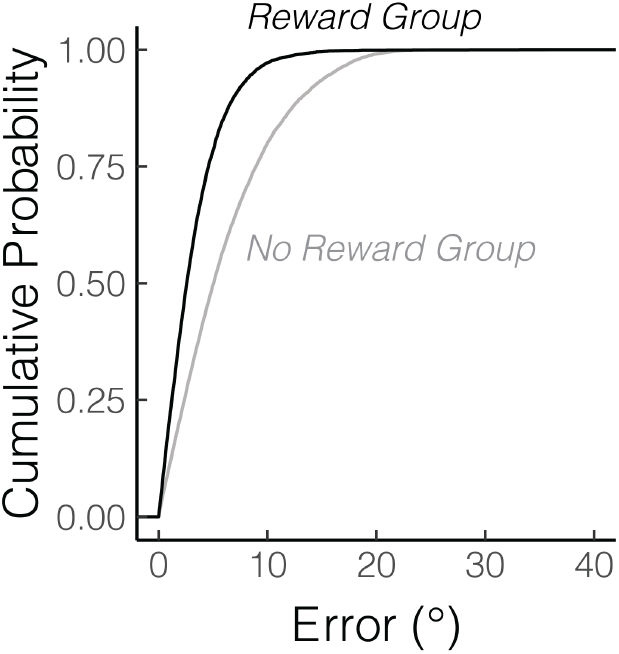
Cumulative probability for context trials as a function of the absolute error.

## Funding

JST is funded by the PODSII scholarship from the Foundation for Physical Therapy Research. RBI is funded by the NIH (NINDS: R35NS116883-01). HEK is funded by NIH K12 (HD055931).

## Notes

### Competing Interest Statement

The authors have declared no competing interest.

